# Identification of peripheral neural circuits that regulate heart rate using optogenetic and viral vector strategies

**DOI:** 10.1101/456483

**Authors:** Pradeep S. Rajendran, Rosemary C. Challis, Charless C. Fowlkes, Peter Hanna, John D. Tompkins, Maria C. Jordan, Sarah Hiyari, Beth A. Gabris-Weber, Alon Greenbaum, Ken Y. Chan, Benjamin E. Deverman, Heike Münzberg, Jeffrey L. Ardell, Guy Salama, Viviana Gradinaru, Kalyanam Shivkumar

## Abstract

Heart rate is under the precise control of the autonomic nervous system. However, the wiring of peripheral neural circuits that regulate heart rate is poorly understood. Here, we developed a clearing-imaging-analysis pipeline to visualize innervation of intact hearts in 3D and employed a multi-technique approach to map parasympathetic and sympathetic neural circuits that control heart rate in mice. We anatomically and functionally identify cholinergic neurons and noradrenergic neurons in an intrinsic cardiac ganglion and the stellate ganglia, respectively, that project to the sinoatrial node. We also report that the heart rate response to optogenetic versus electrical stimulation of the vagus nerve displays different temporal characteristics and that vagal afferents enhance parasympathetic and reduce sympathetic tone to the heart via central mechanisms. Our findings provide new insights into neural regulation of heart rate, and our methodology to study cardiac circuits can be readily used to interrogate neural control of other visceral organs.

---

Situation-dependent changes in heart rate are essential for survival and are under the precise control of the autonomic nervous system (ANS)^1^. Heart rate reduction during sleep^2^ and elevation during exercise^3^ result from changes in parasympathetic and sympathetic tone. In fact, heart rate variability has been utilized extensively as an index of ANS function^4^^-^^6^. Although it is well known that the parasympathetic and sympathetic nervous systems innervate the sinoatrial (SA) node^7^^,^^8^ and regulate heart rate^9^^-^^12^, the wiring of these neural circuits in the periphery is not well characterized. Anatomical and functional maps of these fundamental cardiac circuits are needed to understand physiology, characterize remodeling in disease (e.g., sick sinus syndrome^13^), and develop novel therapeutics. However, these efforts have been hindered by a shortage of tools that target the peripheral nervous system (PNS) with specificity and precision.

Cardiac circuit anatomy has traditionally been studied using thin sections^14^ and whole-mount preparations^15^. However, these methods do not preserve the structure of intact circuits and only provide 2D information. In contrast, tissue clearing methods render tissues optically transparent while preserving their molecular and cellular architecture and can be combined with a variety of labeling strategies to enable 3D visualization of intact circuits^16^^,^^17^. To trace cardiac circuits, dyes and proteins have historically been used. However, achieving cell type-specificity and/or sparse labeling needed for singe-cell tracing and delineating circuit connectivity is difficult or not possible with these methods^18^. Additionally, many peripheral neuronal populations such as intrinsic cardiac ganglia are challenging to access surgically for tracer delivery. Adeno-associated viruses (AAVs) can address these limitations because they can be utilized for efficient *in vivo* gene expression in defined cell populations when used in Cre transgenic animals or with cell-type specific regulatory elements^19^^-^^21^. In addition, intersectional strategies can be used to titrate gene expression to achieve sparse labeling^20^. AAVs can also be delivered systemically to target difficult-to-reach populations^19^^-^^21^.

Functional mapping of cardiac circuits has relied on electrical or pharmacological manipulation of the ANS with simultaneous physiological measurements^22^^,^^23^. However, each of these methods has disadvantages. Electrical techniques lack spatial precision and specificity. Autonomic nerves such as the vagus contain motor and sensory fibers^24^, and electrical stimulation typically activates both fiber types^25^^,^^26^ as well as surrounding tissues^27^. Pharmacological techniques exhibit improved selectivity but lack temporal resolution. In contrast, optogenetics, which uses light-sensitive ion channels (e.g., channelrhodopsin-2 (ChR2), halorhodopsin), enables precise spatiotemporal control of defined cell populations.

Here, we develop a clearing-imaging-analysis pipeline to visualize innervation of whole hearts in 3D and employ a multi-technique approach, which includes AAV-based sparse labeling and tract-tracing, retrograde neuronal tracing with cholera toxin subunit B (CTB), and optogenetics with simultaneous physiological measurements, to map peripheral parasympathetic and sympathetic neural circuits that regulate heart rate in mice.

## RESULTS

### Tissue clearing and computational pipeline to assess global innervation of whole hearts

To characterize global innervation of the mouse heart in 3D, we developed a clearing-imaging-analysis pipeline (Figure 1a). We stained whole hearts with an antibody against the pan-neuronal marker protein gene product 9.5 (PGP9.5) and rendered them optically transparent using an immunolabeling-enabled three-Dimensional Imaging of Solvent-Cleared Organs (iDISCO) protocol (Figure 1b)^28^. We used confocal microscopy to image large tissue volumes (Figure 1c and Supplemental Movie 1) and lightsheet microscopy to image entire hearts (Supplemental Movie 2)^29^^,^^30^. We observed cardiac ganglia surrounding the pulmonary veins and a dense network of nerve fibers coursing through the atrial and ventricular myocardium (Figure 1c). Innervation was seen throughout the entire thickness of the myocardium, with large-diameter nerve fiber bundles located near the epicardium and smaller fiber bundles in the mid-myocardium and endocardium (Figure 1c and Supplemental Movie 3). To analyze these data, we created a semiautomated computational pipeline to detect nerve fibers over large tiled volumes and to measure microanatomical features of fibers such as diameter and orientation (Figure 1, d and e). Large-diameter nerve fiber bundles typically entered near the base of the heart. These bundles coursed perpendicular to the atrioventricular (AV) groove and branched into smaller fiber bundles as they progressed towards the apex. These data from healthy hearts will be important for future characterization of neural remodeling in cardiovascular diseases such as myocardial infarction (MI) in which innervation patterns are disrupted and nerve sprouting occurs^31^^,^^32^.

**Figure 1.**
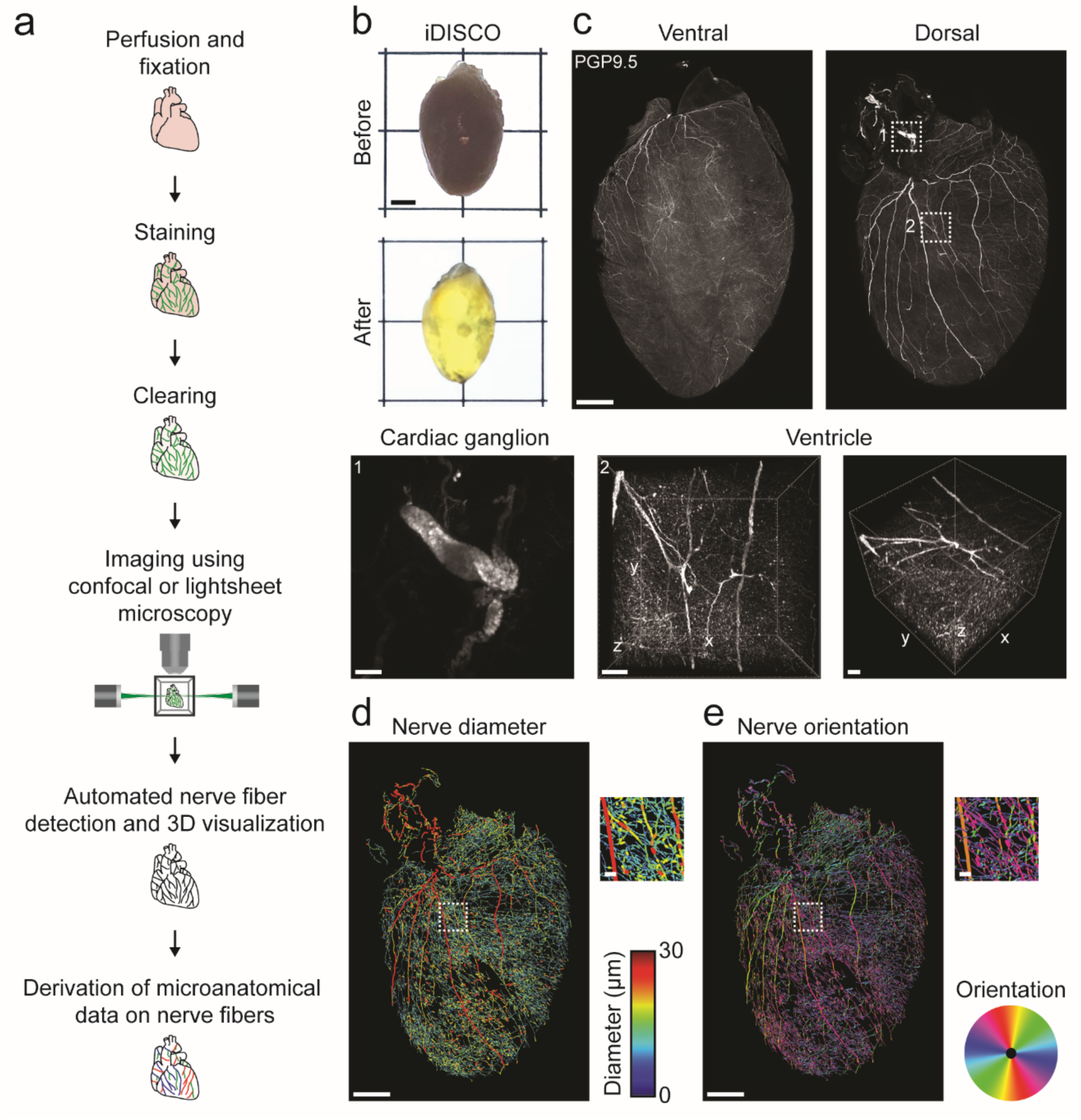
Tissue clearing and computational pipeline to assess global innervation of a whole heart in 3D. (**a**) Schematic of the clearing-imaging-analysis pipeline for assessing cardiac innervation. (**b**) A whole heart (top) was rendered transparent (bottom) using the iDISCO protocol. (**c**) 3D confocal projections of the ventral (1500 µm z-stack) and the dorsal side of cleared heart (1200 µm z-stack) from **a** with PGP9.5 staining (gray). Inset 1 shows a maximum intensity projection (MIP) confocal image of a cardiac ganglion. Inset 2 shows 785 µm-thick 3D projections of the entire left ventricular wall. (**d**, **e**) A semiautomated computational pipeline was used to detect nerve fibers in dorsal heart image from **c** and derive their diameter (**d**) and orientation (**e**). Insets show higher magnification images. Scale bars are 2 mm (**b**), 1 mm (**c** (top), **d** (left), **e** (left)), and 100 µm (**c** (bottom), **d** (right), **e** (right)).

### AAV-based sparse labeling and tract-tracing of cholinergic neurons in intrinsic cardiac ganglia

After visualizing global cardiac innervation, we assessed whether we could identify a subset of cholinergic neurons that form an anatomical circuit with the SA node to potentially regulate heart rate. We used cell type-restricted sparse viral labeling to trace cholinergic neurons in intrinsic cardiac ganglia to their regions of innervation. We systemically co-administered Cre-dependent vectors expressing fluorescent proteins (XFPs) from the tetracycline-responsive element (TRE)-containing promoter at a high dose and the tetracycline transactivator (tTA) from the human synapsin I promoter (hSyn1) at a lower dose in ChAT-IRES-Cre transgenic mice (Figure 2a)^20^. Compared to dense multicolor labeling (Figure 2b), sparse multicolor labeling resulted in a labeling density in intrinsic cardiac ganglia that was lower and would more easily allow for tracing (Figure 2c)^20^. For tract-tracing of cholinergic neurons in intrinsic cardiac ganglia, we utilized sparse single-color labeling with tdTomato. Three weeks after viral delivery, hearts were collected and stained with an antibody for hyperpolarization-activated cyclic nucleotide-gated potassium channel 4 (HCN4) to identify the SA node, the AV node, and the conduction system^33^. We observed cholinergic fibers from a cardiac ganglion, located below the junction of the right and left inferior pulmonary veins (inferior pulmonary veins-ganglionated plexus, IPV-GP), projecting towards the SA node, the AV node, and the ventricles (Figure 2d), identifying cholinergic neurons that are potentially involved in chronotropic, dromotropic, and ventricular control, respectively.

**Figure 2.**
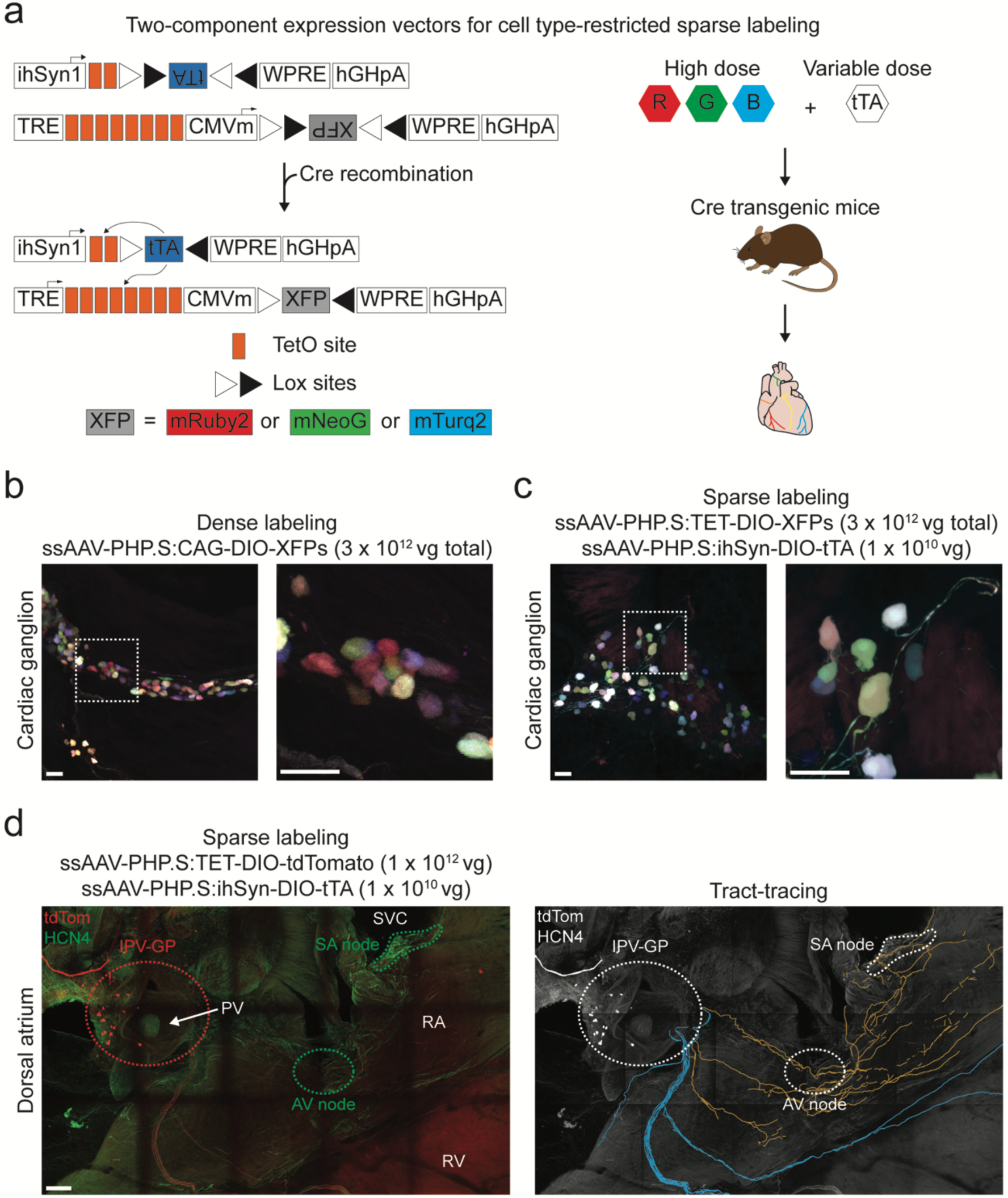
AAV-based sparse labeling and tract-tracing of cholinergic neurons in intrinsic cardiac ganglia. (**a**) Schematic of the two-component expression system for cell type-restricted, sparse multicolor labeling. Expression of XFPs is dependent on cotransduction of an inducer in Cre-expressing cells. The dose of the inducer vector can be titrated to control extent of XFP labeling. (**b**) To densely label cholinergic neurons in cardiac ganglia, ChAT-IRES-Cre mice were systemically co-administered 3 Cre-dependent vectors expressing either mRuby2, mNeonGreen, or mTurquoise2 from the ubiquitous CAG promoter (ssAAV-PHP.S:CAG-DIO-XFP) (1 x 10^12^ vector genomes (vg) each; 3 x 10^12^ vg total). A MIP image of a cardiac ganglion. Inset shows a higher magnification image. (**c**) To sparsely labeling cholinergic neurons in cardiac ganglia, ChATIRES-Cre mice were systemically co-administered 3 Cre-dependent vectors expressing XFPs from the TRE-containing promoter (ssAAV-PHP.S:TRE-DIO-XFP) (1 x 10^12^ vg each; 3 x 10^12^ vg total) and a Cre-dependent inducer vector expressing the tTA from the ihSyn1 promoter (ssAAV-PHP.S:ihSyn1-tTA) (1 x 10^10^ vg) (right). A MIP image of a cardiac ganglion. Inset shows a higher magnification image. (**d**) To trace cholinergic fibers from a cardiac ganglion located below the inferior pulmonary veins (PV) (inferior pulmonary veins-ganglionated plexus, IPV-GP), sparse labeling was performed by systemically co-administering ssAAV-PHP.S:TET-DIO-tdTomato at a high dose (1 x 10^12^ vg) and ssAAV-PHP.S:ihSyn1-DIO-tTA at a lower dose (1 x 10^10^ vg). A MIP image of the dorsal atrium with native tdTomato fluorescence (red) and HCN4 staining (green) (left). Fibers were traced from the IPV-GP with neuTube and overlaid on a grayscale MIP image (right). Orange fibers projected to the right atrium (RA) including the sinoatrial (SA) and atrioventricular (AV) nodes and blue fibers to the ventricles. Scale bars are 50 µm (**b**, **c**) and 200 µm (**d**). All images were acquired on tissue collected 3 weeks after intravenous injection. RV, right ventricle; SVC, superior vena cava.

### *Ex vivo* optogenetic stimulation of cholinergic neurons in the IPV-GP

Next, to functionally assess whether cholinergic neurons in the IPV-GP regulate heart rate, AV conduction, and ventricular electrophysiology, we used an optogenetic approach. We expressed ChR2 in cholinergic neurons by crossing transgenic ChAT-IRES-Cre mice with reporter mice containing a Cre-dependent ChR2-enhanced yellow fluorescent protein allele (ChR2-eYFP; offspring from this cross are subsequently referred to as ChAT-ChR2-eYFP mice). ChR2-eYFP expression in intrinsic cardiac neurons was confirmed by staining hearts for GFP (to amplify ChR2-eYFP detection) and PGP9.5 (Figure 3, a and b). Almost all PGP9.5+ neurons in cardiac ganglia were GFP+ (96.4 ± 1.2%) (Figure 3c). GFP staining was also present in PGP9.5+ nerve fibers in the atria and ventricles (Figure 3b). These data are consistent with previous studies reporting that the majority of intrinsic cardiac neurons are cholinergic^34^ and that the ventricles as well as atria receive cholinergic innervation^34^^-^^36^.

**Figure 3.**
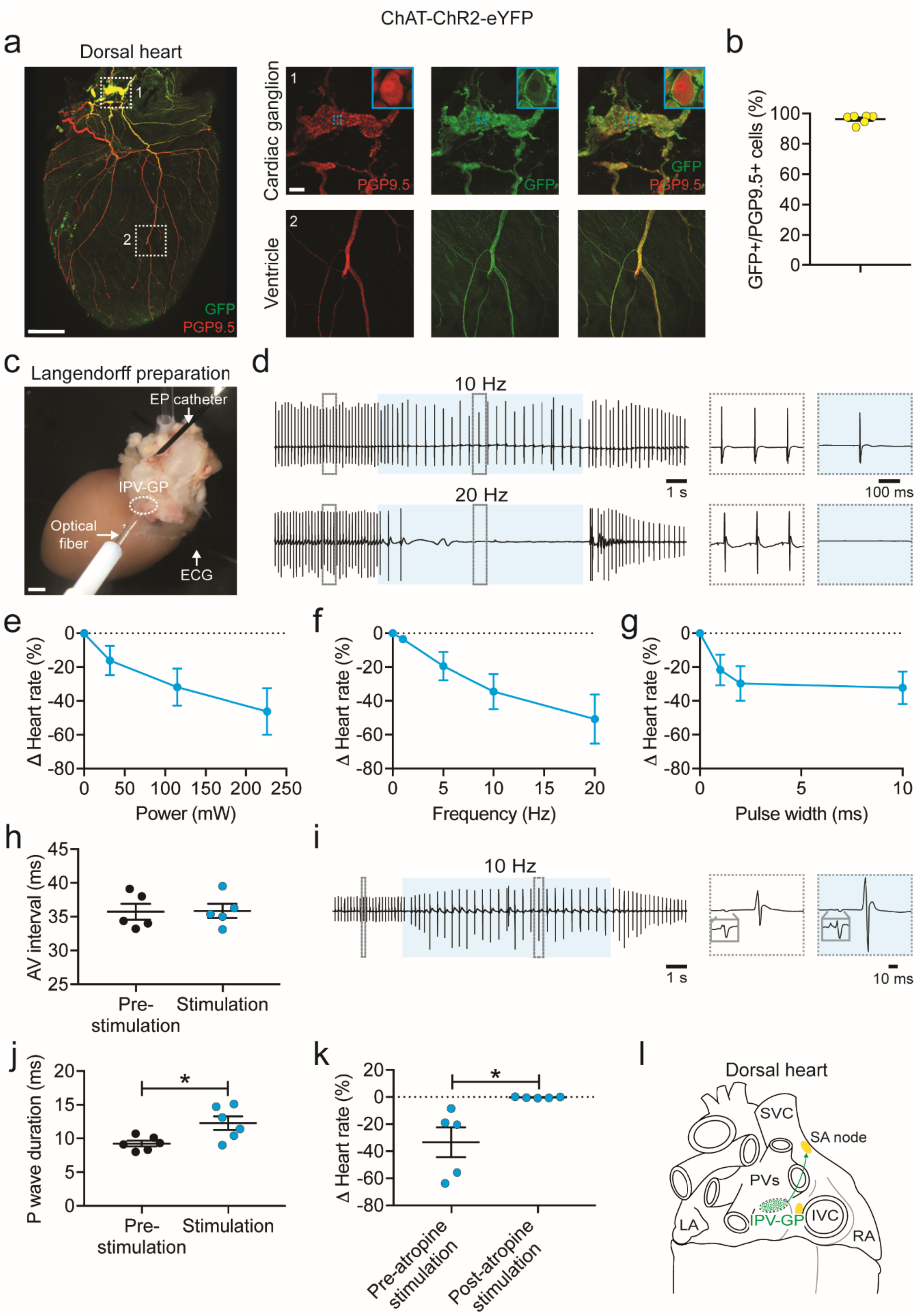
*Ex vivo* optogenetic stimulation of cholinergic neurons in the IPV-GP. (**a**) A 3D projection (1200 µm z-stack) of the dorsal side of a heart from a ChAT-ChR2-eYFP mouse with PGP9.5 (red) and GFP staining (green). Insets 1 and 2 show MIP images of a cardiac ganglion and the ventricle, respectively. Blue dashed boxes indicate location of higher magnification images in blue boxes. (**b**) Percentage of PGP9.5+ cells expressing GFP in cardiac ganglia. (**c**) Langendorff-perfused hearts were used for optogenetic stimulation of cholinergic neurons in a cardiac ganglion. A blue laser-coupled optical fiber was positioned for focal illumination of the IPV-GP (circle). A surface electrocardiogram (ECG) was recorded with bath electrodes and intracardiac electrograms with an electrophysiology (EP) catheter. (**d**) Representative ECGs during stimulation (blue shading) at 10 Hz, 10 ms, and 221 mW (top) and 20 Hz, 10 ms, and 221 mW (bottom). Insets show the ECGs before and during stimulation. (**e**-**g**) Dose response curves summarizing the effects of altering light pulse power (**e**), frequency (**f**), and pulse width (**g**) on heart rate. (**h**) Summary of the AV interval before and during stimulation at 10 Hz and 10 ms (*P* = 0.877). (**i**) Representative ECG during stimulation (blue shading) at 10 Hz, 10 ms, and 221 mW. Insets show a single beat before and during stimulation, with gray boxes showing a higher magnification of the P wave. (**j**) The P wave duration before versus during stimulation (*P =* 0.033). (**k**) Summary of the heart rate response to stimulation before versus after atropine administration (*P* = 0.040). (**l**) Cartoon depicting IPV-GP-SA node circuit. *n =* 6 mice (**b**, **j**) and 5 mice (**e**-**h**, **k**); mean ± s.e.m.; paired, two-tailed *t*-test (\**P* < 0.05). Scale bars are 1 mm (**a** (left), **c**) and 100 µm (**a** (right)). IVC, inferior vena cava; LA, left atrium.

After verifying ChR2 expression, we next assessed whether selective stimulation of cholinergic neurons in the IPV-GP modulated heart rate, AV nodal conduction, and ventricular electrophysiology using optogenetics in *ex vivo* Langendorff-perfused hearts. A blue laser-coupled optical fiber was positioned for focal illumination of the IPV-GP while cardiac electrical activity was simultaneously recorded (Figure 3, c and d). Optogenetic stimulation resulted in a decrease in heart rate that was dependent on light pulse power, frequency, and pulse width (Figure 3, e-g) but did not change the AV interval (35.7 ± 1.2 ms versus 35.9 ± 1.0 ms; *P* = 0.88) (Figure 3h). The lack of change in AV nodal conduction suggests that fibers from this ganglion may pass through the AV node without synapsing. In addition, stimulation prolonged the P wave duration (9.3 ± 1.0 ms versus 12.3 ± 2.4 ms; *P* < 0.05) and caused P wave fractionation (Figure 3, i and j). P wave fractionation has been reported in humans following administration of adenosine ^37^, which mimics the effects of acetylcholine released from cholinergic nerve terminals ^38^^,^^39^. During stimulations at higher frequencies, we occasionally observed ectopic atrial rhythms (*n* = 3/5 mice) and even asystole (*n* = 1/5 mice) (Figure 3d), demonstrating the profound effect of the IPV-GP on the SA node and atrial function. The response to stimulation was abolished by administration of the muscarinic receptor antagonist atropine (−33.5 ± 11% versus −0.3 ± 0.2%; *P* < 0.05) (Figure 3k), indicating that the bradycardic response was indeed mediated by selective stimulation of cholinergic neurons.

Since ChR2 is expressed in both preganglionic cholinergic inputs to and postganglionic cholinergic neurons in the IPV-GP in ChAT-ChR2-eYFP mice, we assessed whether we could stimulate only postganglionic cholinergic neurons in the IPV-GP using optogenetics and still modulate heart rate. We first evaluated whether we could preferentially deliver transgenes to peripheral cholinergic neurons in intrinsic cardiac ganglia rather than central cholinergic neurons in the medulla using systemic AAVs. We used AAV-PHP.S, a capsid variant that more efficiently transduces the PNS and many visceral organs including the heart, as compared to AAV9^20^. We packaged a Cre-dependent genome that expresses eYFP from the ubiquitous CAG promoter into AAV-PHP.S and systemically administered the virus to ChAT-IRES-Cre transgenic mice. Three weeks later, we evaluated eYFP expression in the medulla, the vagus nerve, and intrinsic cardiac ganglia with GFP staining. Central cholinergic neurons in the dorsal motor nucleus of the vagus nerve did not express detectable levels of eYFP and cholinergic fibers in the vagus nerve were weakly transduced (Supplemental Figure 1, a-c). In contrast, we observed robust transduction of peripheral cholinergic neurons in intrinsic cardiac ganglia (80.1 ± 1.5% PGP9.5+ cells expressed GFP) (Supplemental Figure 1, d and e), likely due to the strong tropism AAV-PHP.S displays for the PNS over the CNS. For functional studies, we packaged a Cre-dependent genome that expresses ChR2-eYFP from the ubiquitous CAG promoter in AAV-PHP.S, systemically administered the virus to ChAT-IRES-Cre transgenic mice, and evaluated expression 5 weeks later. In Langendorff-perfused hearts, we were able to optogenetically stimulate postganglionic cholinergic neurons in the IPV-GP and decrease heart rate (Supplemental Figure 2). Taken together, our anatomical and functional data establish an IPV-GP-SA node circuit involved in heart rate regulation (Figure 3j). Furthermore, the engineered AAV, AAV-PHP.S^20^, can be a powerful tool to dissect out the roles of peripheral versus central circuits on organ control.

### *In vivo* optogenetic versus electrical stimulation of the vagus nerve

Electrical vagus nerve stimulation (VNS) has been used in numerous preclinical and clinical studies for the treatment of cardiovascular diseases^40^ and other conditions (e.g., rheumatoid arthritis^41^). However, the relative contributions of vagal efferent and afferent fibers on cardiac function are not well understood because conventional techniques do not allow for fiber type-specific stimulation. To address this limitation, we examined whether we could selectively stimulate efferent fibers in the vagus nerve using optogenetics. We also assessed whether there was a difference between optogenetic and electrical VNS on heart rate (Figure 4a), as a previous study showed that electrical stimulation of motor nerves results in a non-orderly, non-physiological recruitment of fibers, with larger fibers activated first^42^. The vagus and other autonomic nerves contain both motor and sensory fibers that vary in diameter and myelination^27^ and non-electrical techniques such as optogenetics are needed to study their physiological role.

**Figure 4.**
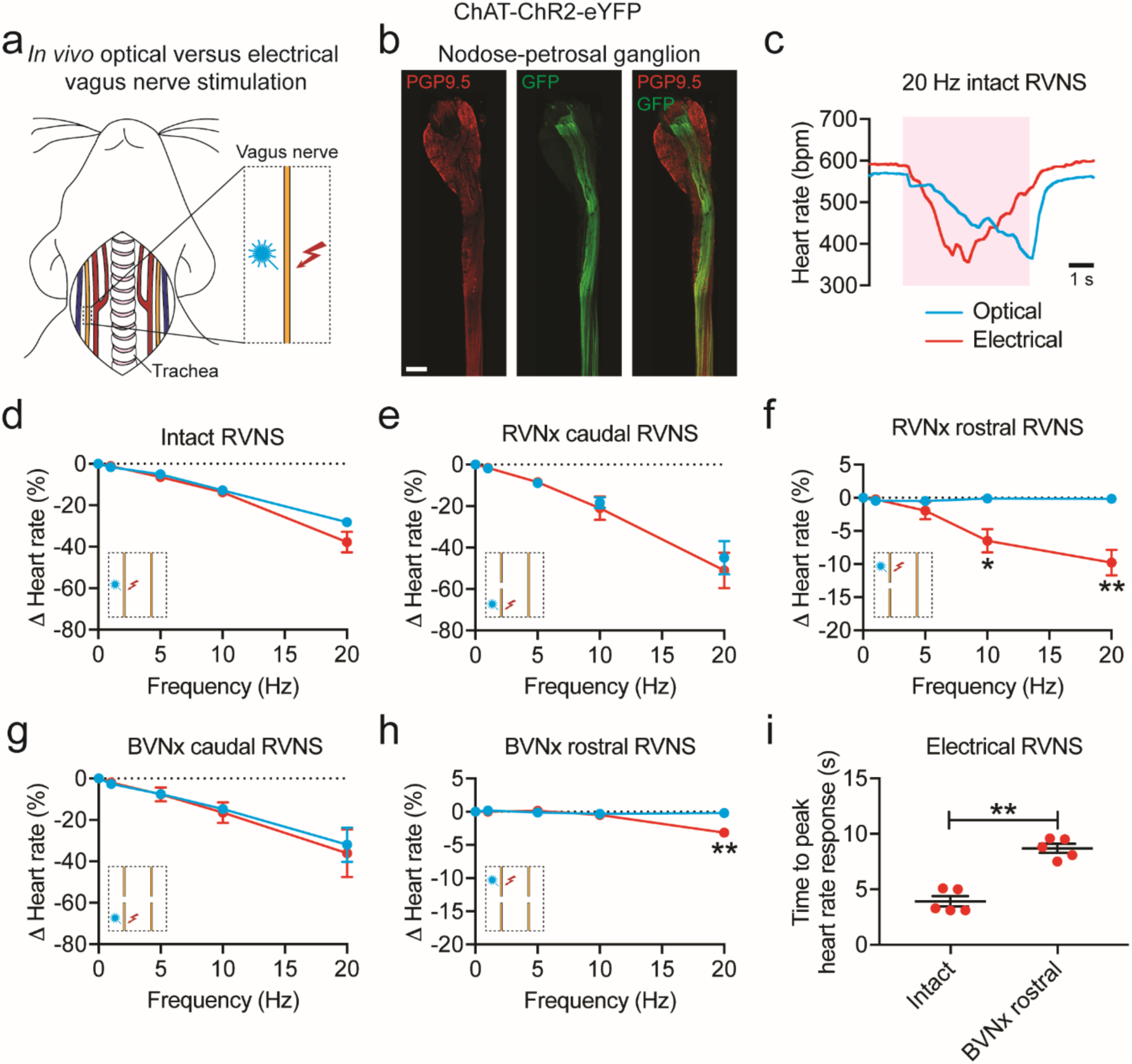
*In vivo* optogenetic versus electrical stimulation of the vagus nerve. (**a**) Cartoon depicting optogenetic and electrical vagus nerve stimulation strategy in ChAT-ChR2-eYFP mice. The right vagus nerve was surgically exposed in anesthetized mice and either light or electricity was used for stimulation. (**b**) MIP images of the right nodose-petrosal ganglion complex and vagus nerve with PGP9.5 (red) and GFP staining (green). (**c**) Representative heart rate responses during optogenetic versus electrical right vagus nerve stimulation (RVNS; magenta shading) at identical frequencies (20 Hz) and pulse widths (10 ms) in the intact state. The light pulse power for optogenetic stimulation was 57 mW and the current for electrical stimulation was 5 µA. (**d**) Frequency response curve summarizing the effects of optogenetic versus electrical RVNS on heart rate in the intact state. (**e**, **g**) Frequency response curve summarizing the effects of optogenetic versus electrical RVNS of the caudal end on heart rate following right vagotomy (RVNx) (**e**) and bilateral vagotomy (BVNx) (**g**). (**f**, **h**) Frequency response curve summarizing the effects of optogenetic versus electrical RVNS of the rostral end on heart rate following RVNx (*P* = 0.023 at 10 Hz; *P* = 0.007 at 20 Hz) (**f**) and BVNx (*P* = 0.001 at 20 Hz) (**h**). In **d**-**h** insets show a schematic of the stimulation protocol. (**i**) The time to peak heart rate response during electrical RVNS in the intact state versus of the rostral end following BVNx (*P* = 0.003). *n* = 8 mice (**d**), 3 mice (**e**, **g**), and 5 mice (**f**, **h**, **i**); mean ± s.e.m.; paired, two-tailed *t*-test (\*\**P* < 0.01; \**P* < 0.05). Scale bar is 200 µm (**b**).

To confirm that ChR2-eYFP expression was limited to vagal efferents in ChAT-ChR2-eYFP mice, we stained for GFP and PGP9.5 in the nodose-petrosal ganglion complex, which contains the cell bodies of vagal sensory neurons, and the cervical vagus nerve (Figure 4b). eYFP was not detected in PGP9.5+ cell bodies in the nodose-petrosal ganglion complex and was only present in a subset of PGP9.5+ vagal fibers (*n =* 5 mice). After verifying expression, we next performed functional studies in anesthetized mice in which we positioned a laser-coupled optical fiber for focal illumination above and a hook electrode underneath the right vagus nerve. In this context, optogenetic VNS activates only efferent fibers (GFP+), whereas electrical VNS presumably activates subsets of efferent and afferent fibers (PGP9.5+) in the vagus nerve (Figure 4b). With both vagi intact, optogenetic and electrical right VNS resulted in a similar decrease in heart rate (Figure 4d); however, the slopes of the responses were dramatically different (*n =* 6/8 mice; Figure 4c), likely due to differential fiber recruitment^42^. The heart rate response to optogenetic versus electrical stimulation of the caudal end of the right vagus nerve was also similar following either right or bilateral vagotomy (Figure 4, e and g). In contrast, optogenetic stimulation of the rostral end of the right vagus nerve following either right or bilateral vagotomy did not affect heart rate, whereas electrical stimulation surprisingly resulted in a decrease in heart rate at 10 Hz (−6.5 ± 1.8% versus −0.1 ± 0.1% following right vagotomy for electrical versus optogenetic stimulation; *P* < 0.05) and 20 Hz (−9.8 ± 1.9% versus −0.2 ± 0.1% following right vagotomy and −3.2 ± 0.4% versus −0.2 ± 0.1% following bilateral vagotomy for electrical versus optogenetic stimulation; *P* < 0.01 for both) (Figure 4, f and h). The decrease in heart rate to electrical stimulation of the rostral end of the right vagus nerve following right vagotomy (Figure 4f, red line) was greater than that following bilateral vagotomy (Figure 4h, red line) (*P* < 0.035), suggesting that the response following right vagotomy was in part due to vagal afferent-mediated increase in parasympathetic efferent outflow through the intact contralateral vagus nerve. In addition, there was an increased latency to peak heart rate response with electrical stimulation of the rostral right vagus nerve following bilateral vagotomy compared to that of the intact right vagus nerve (8.7 ± 0.4 ms versus 3.9 ± 0.5 ms; *P* < 0.01) (Figure 4i), indicating that the response post-bilateral vagotomy was likely due to vagal afferent-mediated withdrawal of sympathetic tone^43^. Taken together, our data suggest that optogenetic stimulation selectively activates vagal efferents in ChAT-ChR2-eYFP mice and that vagal afferents act centrally to 1) increase parasympathetic efferent outflow and 2) decrease sympathetic efferent outflow to the heart.

### Localization and *in vivo* optogenetic stimulation of cardiac-projecting noradrenergic neurons in the stellate ganglia

The sympathetic nervous system, along with the parasympathetic nervous system, precisely regulates heart rate in normal physiology. To anatomically and functionally dissect noradrenergic neurons that form a circuit with the SA node, we used a retrograde neuronal tracer and an optogenetic approach. Noradrenergic nerve fibers densely innervate the heart as shown by staining for tyrosine hydroxylase (TH), the rate-limiting enzyme in norepinephrine synthesis (Figure 5a). To identify the location of cardiac-projecting sympathetic neurons, we injected the retrograde neuronal tracer CTB conjugated to Alexa Fluor 488 into the heart (Figure 5b). The majority of CTB+ neurons were located in the stellate ganglia of the paravertebral chain, with fewer labeled neurons in the middle cervical and second thoracic (T2) ganglia (Figure 5c). On average, 236 ± 39 neurons were labeled in the right and 261 ± 34 neurons labeled in the left stellate ganglion (*n* = 5 mice). Heat maps of the right and left stellate ganglia show that cardiac-projecting sympathetic neurons are clustered in the craniomedial aspect (Figure 5d) and suggesting that these ganglia may have a viscerotopic organization.

**Figure 5.**
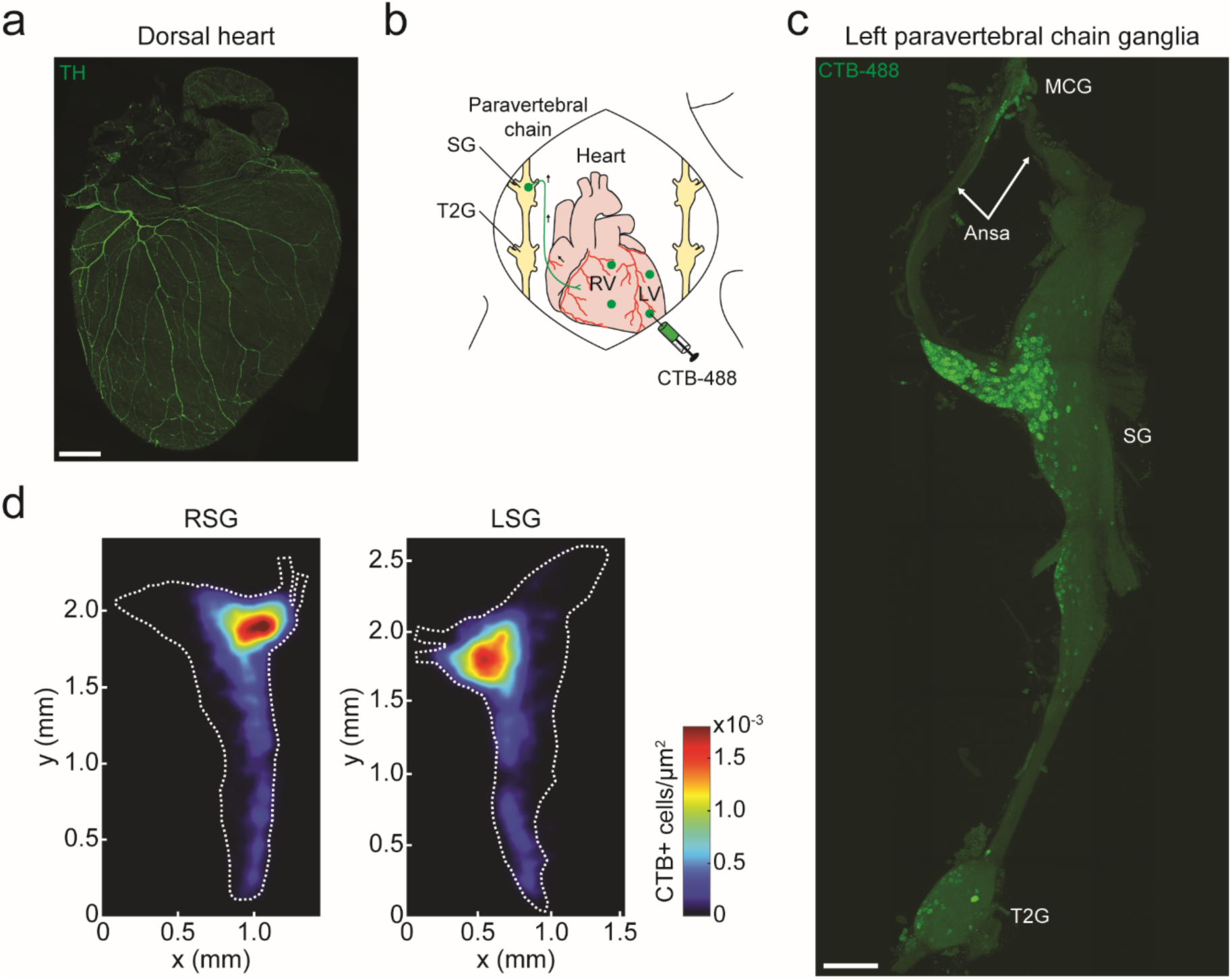
Cardiac-projecting neurons in stellate ganglia (SG) are clustered in craniomedial aspect. (**a**) A 3D projection (1200 µm z-stack) of the dorsal side of a C57BL/6J mouse heart with TH staining (green). (**b**) Cartoon depicting CTB-Alexa Fluor 488 (CTB-488) injections into the heart to retrogradely trace neurons in the paravertebral chain ganglia that project to the heart. (**c**) A MIP image of the left paravertebral chain from the middle cervical ganglion (MCG) to the second thoracic ganglion (T2G) showing the location of CTB-488+ neurons that project to the heart. (**d**) Summary heat map of the right stellate ganglion (RSG) and the left stellate ganglion (LSG). *n* = 5 mice (**d**). Scale bars are 1 mm (**a**, **c**). CTB injections were performed in C57BL/6J mice. LV, left ventricle.

Next, we assessed whether we could selectively stimulate noradrenergic neurons in the paravertebral chain using optogenetics and modulate heart rate. We expressed ChR2 in noradrenergic neurons by crossing transgenic TH-IRES-Cre mice with reporter mice containing a Cre-dependent ChR2-eYFP allele (offspring from this cross are subsequently referred to as TH-ChR2-eYFP mice). ChR2-eYFP expression in stellate ganglion neurons was confirmed by staining for GFP and TH (Figure 6b). Almost all TH+ neurons were also GFP+ (95.9 ± 1.6%; *n* = 4 mice). After verifying expression, we performed functional studies in open-chest anesthetized mice in which we positioned a laser-coupled optical fiber for focal illumination above the craniomedial right stellate ganglion (RSG) or right T2 ganglion (Figure 6a). Optogenetic stimulation of the craniomedial RSG resulted in a frequency- and pulse width-dependent increase in heart rate (Figure 6, d and e). Although a small number of cardiac-projecting neurons were located in the T2 ganglion (Figure 5c), there was no heart rate response to stimulation of this ganglion (Figure 6f). The response to craniomedial RSG stimulation was abolished by administration of the β-adrenergic receptor antagonist propranolol (5.6 ± 1.4% versus 0.2 ± 0.1%; *P* < 0.05) (Figure 6g), indicating that the tachycardic response was indeed mediated by selective stimulation of noradrenergic neurons. Taken together, our anatomical and functional data establish a craniomedial RSG-SA node circuit involved in heart rate regulation.

**Figure 6.**
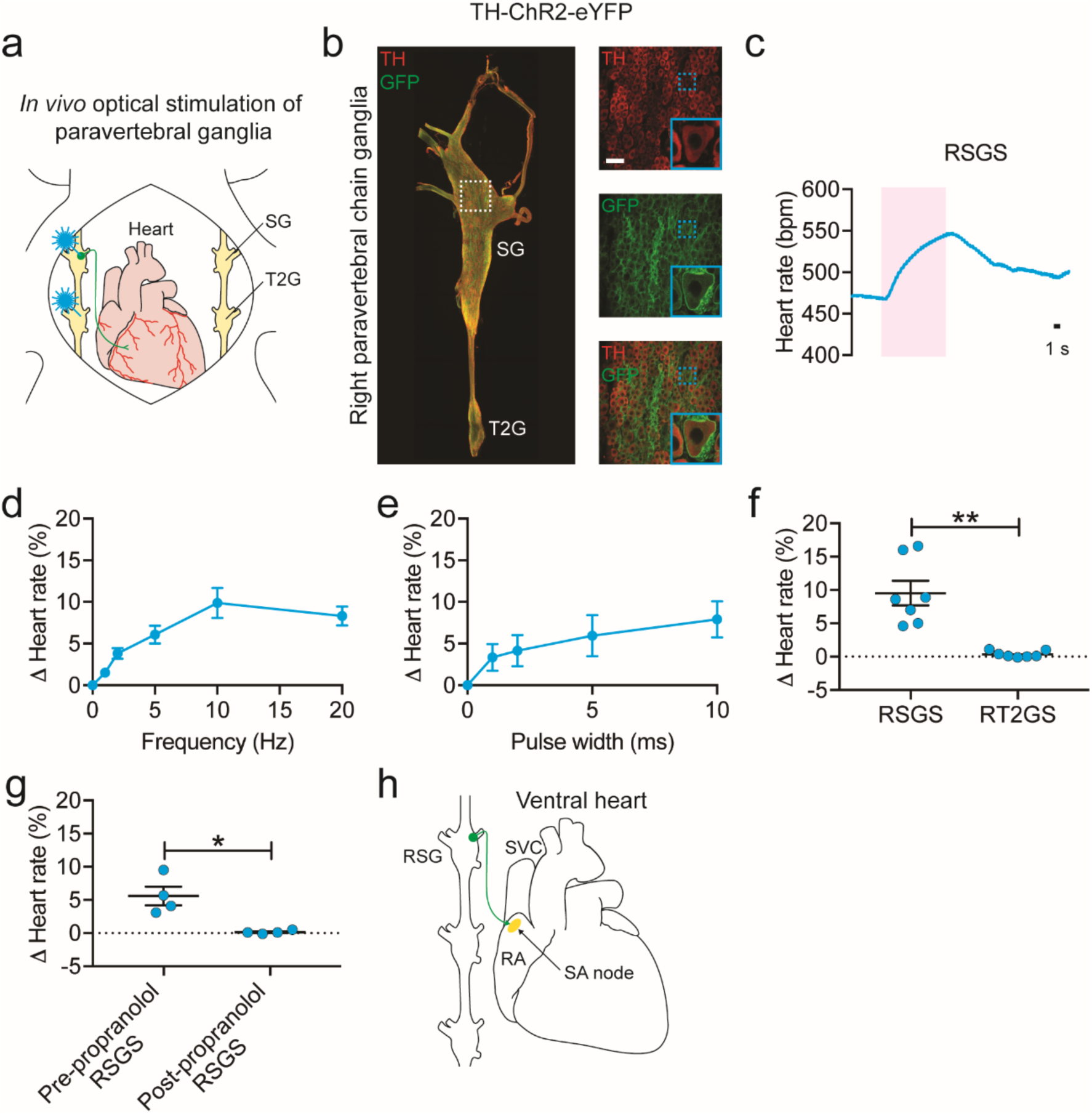
*In vivo* optogenetic stimulation of noradrenergic neurons in the SG. (a) Cartoon depicting optogenetic SG and T2G stimulation strategy in TH-ChR2-eYFP mice. The right paravertebral chain was surgically exposed in anesthetized mice and light was used for stimulation. (**b**) A MIP image of the right paravertebral chain showing the SG and the T2G with TH (red) and GFP staining (green). Inset shows single-plane images of SG. Blue dashed boxes indicate location of higher magnification images in blue boxes. (**c**) Representative heart rate response during 10 Hz, 10 ms, and 126 mW craniomedial RSG stimulation (RSGS). (**d**, **e**) Dose response curves summarizing the effects of altering craniomedial RSGS frequency (**d**) and pulse width (**e**) on heart rate. (**f**) Summary of the heart rate response to craniomedial RSGS versus right T2G stimulation (RT2GS) (*P* = 0.001). (**g**) Summary of the heart rate response to craniomedial RSGS before versus after propranolol administration (*P* = 0.029). (**h**) Cartoon depicting craniomedial RSG-SA node circuit*. n* = 7 mice (**d**, **f**), 5 mice (**e**), and 4 mice (**g**); mean ± s.e.m.; paired, two-tailed *t*-test (\*\**P* < 0.01; \**P* < 0.05). Scale bars are 200 µm (**b** (left) and 50 µm (**b** (right).

## DISCUSSION

We developed a clearing-imaging-analysis pipeline to visualize innervation of whole hearts and employed a multi-technique approach to dissect fundamental parasympathetic and sympathetic neural circuits involved in heart rate regulation. We report several novel findings: 1) cholinergic neurons in the IPV-GP and noradrenergic neurons in the craniomedial RSG project to the SA node and modulate its function; 2) the evoked cardiac response to optogenetic versus electrical stimulation of the vagus nerve displays different temporal characteristics; and 3) vagal afferents enhance parasympathetic and reduce sympathetic efferent outflow to the heart via central mechanisms.

Despite advances in tissue clearing^32^^,^^44^ and imaging techniques^29^^,^^30^, high-resolution, 3D datasets of global cardiac innervation do not exist. We show, for the first time, innervation of an entire cleared mouse heart with cardiac ganglia located around the pulmonary veins and a dense network of nerve fibers throughout the myocardium. To analyze these data, we developed a semiautomated computational pipeline to detect nerve fibers and to measure microanatomical features such as diameter and orientation. These analytical tools and resulting measurements are needed to build a reference atlas of cardiac innervation and for quantitative descriptions of innervation in healthy versus diseased states such as MI. Following MI, innervation around the infarct scar and of remote regions of the heart is altered^31^^,^^32^^,^^45^, and this neural remodeling can modulate the arrhythmia substrate^46^. Understanding changes in innervation post-MI can provide new insights into arrhythmia mechanisms. Furthermore, our clearing-imaging-analysis pipeline can be readily applied to assess innervation of other visceral organs.

It is well known that the parasympathetic and sympathetic nervous systems are critical for heart rate regulation. The SA node and conduction system are densely innervated^7^^,^^8^, and stimulation of the vagus nerve^9^, stellate ganglia^10^, and noradrenergic fibers^11^^,^^12^ modulates heart rate. However, the precise wiring of the underlying neural circuits has not been delineated. We used a novel sparse AAV labeling system^20^ and an optogenetic approach to anatomically and functionally characterize cholinergic neurons that regulate heart rate. We identified cholinergic neurons in the IPV-GP that project to the SA node, the AV node, and the ventricles. Selective optogenetic stimulation of cholinergic neurons in the IPV-GP modulated heart rate and ventricular electrophysiology but, interestingly, not AV nodal conduction. Previous studies showed that electrical stimulation of pulmonary vein ganglia results in biphasic heart rate responses (initial bradycardia followed by tachycardia)^22^^,^^23^. However, since electrical stimulation is non-specific, it is difficult to interpret whether the biphasic response was due to activation of a mixed population of neurons contained in intrinsic cardiac ganglia (i.e., parasympathetic, sympathetic, and sensory)^14^ and/or pass through fibers. Our findings demonstrate an IPV-GP-SA node circuit and highlight the importance of using techniques such as optogenetics, which confer cell type-specificity, to dissect cardiac neural circuity. Furthermore, electrical and optogenetic techniques (using traditional transgenic and AAV-based approaches for ChR2 delivery) stimulate both central preganglionic inputs to and postganglionic neurons in intrinsic cardiac ganglia. Therefore, we used a novel engineered AAV, AAV-PHP.S^20^, that has a strong tropism for the PNS to preferentially deliver ChR2 to postganglionic cholinergic neurons on the heart rather than preganglionic cholinergic neurons in the medulla. Future studies of peripheral neural circuits should use AAV-PHP.S and other engineered AAVs^19^^,^^21^ to dissect the role of central versus peripheral neuronal populations on organ function.

To map noradrenergic neurons that regulate heart rate, we used a retrograde neuronal tracer and optogenetic approach. A previous study in canines using horseradish peroxidase showed that sympathetic postganglionic neurons that innervate the heart are primarily located in the middle cervical ganglia of the paravertebral chain^47^. However, we report, using CTB and confirm with optogenetic stimulation, that the stellate ganglia have a viscerotopic organization with cardiac-projecting neurons clustered in the craniomedial aspect, consistent with a study in cats^48^. In addition to the heart, the stellate ganglia project to many other thoracic structures, including the sweat glands in the forepaw^49^, the lung and trachea^50^, the esophagus^48^, and brown fat^51^. Characterizing stellate ganglia target innervation and cell-type specification is an area of ongoing investigation^52^^,^^53^ that is of interest from a developmental, physiological, and therapeutic perspective.

Current understanding of the role of vagal efferent and afferent fibers on cardiac function is largely based on studies using electrical stimulation, which is non-specific and results in non-orderly, non-physiological recruitment of fibers^42^. Electrical stimulation of the vagus nerve typically activates large-diameter myelinated A fibers, followed by medium-diameter myelinated B fibers and then small-diameter unmyelinated C fibers^27^. *In vivo* we report that optogenetic stimulation of motor fibers in the vagus nerve results in a heart rate response that has a slower onset than electrical stimulation of motor and sensory fibers, likely due to differential fiber recruitment^42^. Our findings suggest that non-electrical techniques such as optogenetics are needed to characterize neural control of cardiac physiology. We also show that activation of vagal afferents decreases heart rate by enhancing parasympathetic efferent outflow and reducing sympathetic efferent outflow centrally, consistent with a prior study showing that global activation of vagal afferents in Vglut2-ChR2 mice and selective activation of Npy2r-ChR2 vagal afferents causes a profound bradycardia^54^. In support of our functional data, anatomical tracing studies have previously shown that cardiac vagal afferent neurons in the nodose-petrosal ganglion complex project to neurons in the nucleus tractus solitarii of the medulla^55^. These neurons then project to the nucleus ambiguus and the dorsal motor nucleus of the vagus nerve in the medulla to modulate parasympathetic efferent outflow^55^ and to the paraventricular nucleus of the hypothalamus to modulate sympathetic efferent outflow^56^. Future studies aimed at identifying cardiac-specific vagal efferent and afferent fibers, similar to those performed in the lungs^54^ and the gastrointestinal system^57^, are needed to better understand vagal control of cardiac physiology and to design next-generation VNS therapies.

Overall, our data highlight the complexity of cardiac neural circuitry and demonstrate that a multi-technique approach is needed to delineate circuit wiring. Understanding the neural control of organ function in greater detail is critical as neuromodulation therapies are emerging as promising approaches to treat a wide range of diseases. Tools such as optogenetics and AAVs are already providing new scientific insights into the structure and function of peripheral neural circuits^54^^,^^57^. A combination of these approaches will help disentangle neural control of autonomic physiology and enable a new era of targeted neuromodulation approaches.

## METHODS

### Animals

All animal experiments were conducted in accordance with the NIH Guide for the Care and Use of Laboratory Animals and were approved by the UCLA and California Institute of Technology Institutional Animal Care and Use Committee. ChAT-IRES-Cre (028861)^58^, Ai32 (024109)^59^, and C57BL/6J mice (000664) were purchased from the Jackson Laboratory. TH-IRES-Cre mice (254)^60^ were purchased from the European Mutant Mouse Archive. ChAT-ChR2-eYFP and TH-ChR2-eYFP mice were created by crossing Ai32 mice with ChAT-IRES-Cre or TH-IRES-Cre mice, respectively. All animals were kept on a 12 h light/dark cycle with ad libitum access to food and water. Data for all experiments were collected from male and female adult mice (greater than 8 weeks old).

### Plasmids

Plasmids used for AAV production include pUCmini-iCAP-PHP.S (Addgene, 103006), pAAVCAG-DIO-mRuby2 (Addgene, 104058), pAAV-CAG-DIO-mNeonGreen^20^, pAAV-CAG-DIO-mTurquoise2 (Addgene, 104059), pAAV-ihSyn1-DIO-tTA (Addgene, 99121), pAAV-TRE-DIO-mRuby2 (Addgene, 99117), pAAV-TRE-DIO-mNeonGreen^20^, pAAV-TRE-DIO-mTurquoise2 (Addgene, 99115), pAAV-TRE-DIO-tdTomato (Addgene, 99116), and pAAV-CAG-DIO-eYFP (Addgene, 104052). pAAV-CAG-DIO-ChR2(H134R)-eYFP was generated by replacing the Ef1a promoter in pAAV-Ef1a-DIO-ChR2(H134R)-eYFP (a gift from Karl Deisseroth, Addgene, 20298) with the CAG promoter from pAAV-CAG-mNeonGreen^20^. The pHelper plasmid was obtained from Agilent’s AAV helper-free kit (Agilent, 240071). CAG, Cytomegalovirus early enhancer element chicken β-actin promoter; DIO = double-floxed inverted open reading frame.

### Virus production and purification

AAVs were produced and purified as previously described^61^. Briefly, AAVs were generated by triple transient transfection of HEK293T cells (ATTC, CRL-3216) using polyethylenimine (Polysciences, 23966-2)^62^. Viral particles were harvested from the media at 72 h post-transfection and from the cells and media at 120 h post-transfection. Virus from the media was precipitated at 4°C with 40% polyethylene glycol (Sigma-Aldrich, 89510)^63^ in 2.5 M NaCl and combined with cell pellets suspended at 37°C in 500 mM NaCl, 40 mM Tris, 10 mM MgCl_2_, and 100 U/ml of salt-active nuclease (ArcticZymes, 70910-202). Clarified lysates were purified over iodixanol (Cosmo Bio USA, OptiPrep, AXS-1114542) step gradients (15%, 25%, 40%, and 60%)^64^. Purified viruses were concentrated, washed in sterile phosphate buffered saline (PBS), sterile filtered, and tittered using quantitative PCR^65^.

### Systemic delivery of viruses

Intravenous administration of AAV vectors was performed by retro-orbital injection with a 31-gauge needle in 6-8 week old mice as previously described^61^. Following injection, 1-2 drops of proparacaine (Akorn Pharmaceuticals, 17478-263-12) were applied to the cornea to provide local analgesia.

### Optogenetic stimulation and physiological measurements

#### Langendorff-perfused heart and cardiac ganglion stimulation

Mice were given heparin (100U, i.p.) to prevent blood clotting and euthanized with sodium pentobarbital (150 mg/kg, i.p.). Once all reflexes subsided and following a midsternal incision, hearts were rapidly excised and Langendorff-perfused with 37°C modified Tyrode’s solution containing the following (in mM): 112 NaCl, 1.8 CaCl_2_, 5 KCl, 1.2 MgSO_4_, 1 KH_2_PO_4_, 25 NaHCO_3_, and 50 D-glucose at pH 7.4. The solution was continuously bubbled with 95% O_2_ and 5% CO_2_. Flow rate was adjusted to maintain a perfusion pressure of 60-80 mmHg. An optical fiber (400 µm core, Doric Lens) coupled to a diode-pumped solid-state laser light source (473 nm, 400 mW, Optic Engine) was positioned for focal illumination of the IPV-GP under the guidance of a fluorescent stereomicroscope (Leica, M205 FA) fitted with a 1x Plan-Apochromat. An octapolar electrophysiology catheter (1.1F, Transonic) was inserted into the left atrium and ventricle via a small incision in the left atrium and 2 platinum electrodes were positioned in the bath to obtain intracardiac electrograms and a bath electrocardiogram, respectively. Intracardiac electrograms and electrocardiogram were amplified with a differential alternating current amplifier (A-M Systems, Model 1700 and Grass, P511, respectively) and continuously acquired (AD Instruments, PowerLab 8/35). Dose response curves were performed to evaluate the effects of altering light pulse power (at 10 Hz and 10 ms), frequency (at 10 ms and 221 mW), and pulse width (at 10 Hz and 221 mW) on heart rate and AV interval. At the end of the experiment, atropine (10 uM) was administered into the perfusate and stimulation (10 Hz, 10 ms, and 221 mW) was repeated. All stimulations were performed for 10 s with 5 min between stimulations for heart rate to return to baseline values.

#### In vivo vagus nerve and paravertebral ganglia stimulation

Mice were anesthetized with isoflurane (induction at 3-5%, maintenance at 1-3% vol/vol, inhalation), intubated, and mechanically ventilated (CWE, SAR-830). Core body temperature was measured and maintained at 37°C. For VNS, a midline neck incision was performed and the left and right cervical vagus nerves were exposed. A laser-coupled optical fiber was positioned above and bipolar platinum hook electrodes coupled to a constant current stimulator (Grass, PSIU6 and Model S88) below the right vagus nerve for focal optical or electrical stimulation, respectively. Irradiance and current were titrated to achieve a 10% decrease in heart rate with both vagi intact at 10 Hz and 10 ms, which was defined as threshold intensity. Threshold intensity for optogenetic stimulation was 77 ± 6 mW and for electrical stimulation was 33 ± 13 µA. Frequency response curves were performed at threshold intensity and 10 ms. All stimulations were performed for 5 s with 5 min between stimulations for heart rate to return to baseline values. For paravertebral ganglia stimulation, a right lateral thoracotomy was performed at the second intercostal space and the paravertebral chain from the stellate to T2 ganglion was exposed. A laser-coupled optical fiber was positioned for focal illumination above the craniomedial RSG or right T2 ganglion. Dose response curves were performed to evaluate the effect of altering RSG stimulation frequency (at 10 ms and 126 mW) and pulse width (at 10 Hz and 126 mW) on heart rate. The effect of RSG versus right T2 ganglion stimulation (10 Hz, 10 ms, and 126 mW) on heart rate was also evaluated. At the end of the experiment, propranolol (2 mg/kg, i.v.) was administered via a femoral vein and craniomedial RSG stimulation (10 Hz, 10 ms, and 126 mW) was repeated. All stimulations were performed for 10 s with 5 min between stimulations for heart rate to return to baseline values. For vagus nerve and paravertebral ganglia stimulation experiments, a lead II electrocardiogram was obtained by 2 needle electrodes inserted subcutaneously into the right forepaw and the left hindpaw.

#### Electrophysiology data analysis

A minimum of 3 beats were averaged at baseline and during stimulation for all electrophysiological data.

### Cholera toxin subunit B heart injections

Mice were given carprofen (5 mg/kp, s.c.) and buprenorphine (0.05 mg/kg, s.c.) 1 h before surgery. Animals were anesthetized with isoflurane (induction at 5%, maintenance at 1-3%, inhalation), intubated, and mechanically ventilated. Core body temperature was measured and maintained at 37°C. The surgical incision site was cleaned 3 times with 10% povidone iodine and 70% ethanol in H_2_O (vol/vol). A left lateral thoracotomy was performed at the 4th intercostal space, the pericardium opened, and the heart was exposed. Ten microliters of CTB conjugated to Alexa Fluor 488 (0.1% in 0.01 M PBS (vol/vol), ThermoFischer Scientific, C22841) was subepicardially injected in the heart with a 31-gauge needle. The surgical wounds were closed with 6-0 sutures. Buprenorphine (0.05 mg/kg, s.c.) was administered once daily for up to 2 d after surgery. Animals were sacrificed 6 d later for tissue harvest.

### Tissue preparation, immunohistochemistry, and imaging

#### Transcardial perfusion

Mice were given heparin (100U, i.p.) to prevent blood clotting and euthanized with sodium pentobarbital (150 mg/kg, i.p.). Once all reflexes subsided, a midsternal incision was made and animals were transcardially perfused with 50 mL ice-cold 0.01 M PBS containing 100U heparin followed by 50 mL freshly prepared, ice-cold 4% paraformaldehyde (PFA; EMS, RT 15714) in PBS. Tissues were postfixed in 4% PFA overnight at 4°C, washed, and stored in PBS with 0.01 % sodium azide. Langendorff-perfused hearts were immersion fixed in 4% PFA overnight at 4°C at the end of the functional experiments

#### Heart clearing

Whole mouse hearts were stained and cleared using a modified iDISCO protocol^28^. Fixed hearts were dehydrated with a graded methanol series (20%, 40%, 60%, and 80% methanol in H_2_O (vol/vol), each for 1 h at room temperature), washed twice with 100% methanol for 1 h at room temperature, and chilled at 4°C. Hearts were then incubated in 66% dichloromethane/33% methanol overnight at room temperature with agitation, washed twice in 100% methanol for 1 h at room temperature, and chilled to 4°C. Next, hearts were bleached with 5% H_2_O_2_ in methanol (vol/vol) overnight at 4°C. After bleaching, hearts were rehydrated with a graded methanol series, followed by one wash with 0.01 M PBS and 2 washes with 0.01 M PBS with 0.2% Triton X-100, each for 1 h at room temperature. For staining, hearts were permeabilized with 0.01 M PBS with 0.2% Triton X-100, 20% DMSO, and 0.3 M glycine and blocked with 0.01 M PBS with 0.2% Triton X-100, 10% DMSO, and 5% normal donkey serum (NDS), each for 2 d at 37°C with agitation. Hearts were incubated in primary antibody rabbit anti-PGP9.5 (Abcam, ab108986, 1:200) diluted in 0.01 M PBS with 0.2% Tween-20 and 10 mg/ml heparin (PTwH) for 1 week at 37°C with agitation. Hearts were then washed several times in PTwH overnight at room temperature before secondary antibody donkey anti-sheep Cy3 (Jackson ImmunoResearch, 713-165-003, 1:300) incubation in PTwH for 1 week at 37°C with agitation. Primary and secondary Ab were replenished half way through staining. Hearts were then washed several times in PTwH overnight at room temperature. For clearing, stained hearts were dehydrated with a graded methanol series and incubated in 66% dichloromethane/33% methanol for 3 h at room temperature with agitation. Hearts were then washed twice in 100% dichloromethane for 15 min at room temperature. Hearts were stored and imaged in benzyl ether (Millipore Sigma, 108014 ALDRICH; refractive index: 1.55).

#### Immunohistochemistry

Fixed hearts and ganglia were blocked in 0.01 M PBS with 10% NDS and 0.2% Triton X-100 PBS for 6 h at room temperature with agitation. Tissues were then incubated in primary Ab diluted in 0.01 M PBS with 0.2% Triton X-100 and 0.01% sodium azide for 3 nights at room temperature with agitation. The following primary antibodies were used: rabbit anti-HCN4 (Alomone Labs, APC-052, 1:100), rabbit anti-PGP9.5 (Abcam, ab108986, 1:500), chicken anti-GFP (Aves, GFP-1020, 1:1000), and sheep anti-TH (Millipore Sigma, AB1542, 1:200). Tissues were washed several times in 0.01 M PBS overnight before incubation in secondary Ab diluted in 0.01 M PBS with 0.2% Triton X-100 and 0.01% sodium azide for 2 nights at room temperature with agitation. The following secondary antibodies were used: donkey anti-rabbit Cy3 (Jackson ImmunoResearch, 711-165-152, 1:400), donkey anti-chicken 647 (Jackson ImmunoResearch, 703-605-155, 1:400), and donkey anti-sheep Cy3 (Jackson ImmunoResearch, 713-165-003, 1:400). Tissues were washed several times in 0.01 M PBS overnight before being mounted on microscope slides in refractive index matching solution (refractive index = 1.46) ^29^.

Fixed brains were cryopreserved in a 30% sucrose PBS solution for at least 48 h at 4°C. Brains were then embedded in OCT (Tissue-Tek), frozen on dry ice, and stored in −80°C until sectioning. For cryosectioning, 40 µm coronal sections were cut using a cryostat microtome (Leica). Sections of the medulla containing the dorsal motor nucleus of the vagus nerve (Allen Mouse Brain Reference Atlas) were blocked in 0.01 M PBS with 10% NDS and 0.2% Triton X-100 PBS for 1 h at room temperature with agitation. Sections were then incubated in primary Ab diluted in 0.01 M PBS with 10% NDS, 0.2% Triton X-100, and 0.01% sodium azide for 1 night at room temperature with agitation. The following primary antibodies were used: rabbit anti-ChAT (Millipore Sigma, AB143, 1:200) and chicken anti-GFP (Aves Labs, 1:1000). Tissues were washed several times in 0.01 M PBS with 0.2% Triton X-100 before incubation with secondary Ab diluted in 0.01 M PBS with 10% NDS, 0.2% Triton X-100, and 0.01% sodium azide for 2 h at room temperature with agitation. The following secondary antibodies were used: donkey anti-rabbit Cy3 (Jackson ImmunoResearch, 711-165-152, 1:1000) and donkey anti-chicken 647 (Jackson ImmunoResearch, 703-605-155, 1:400). Tissues were washed several times in 0.01 M PBS with 0.2% Triton X-100 before being mounted on microscope slides in Fluoromount-G (Thermo Fisher Scientific, 00-4958-02).

#### Imaging

Images were acquired on a confocal laser scanning microscope (Zeiss, LSM 880) fitted with the following objectives: Fluar 5x/0.25 M27 Plan-Apochromat, 10x/0.45 M27 (working distance 2.0 mm), Plan-Apochromat 25x/0.8 Imm Corr DIC M27 multi-immersion, and LD CApochromat 40x/1.1 W Korr. Cleared heart in Supplementary Video 3 was imaged on a custom-built lightsheet fluorescence microscope fitted with a 10x/0.6 CLARITY objective (Olympus, XLPLN10XSVMP) as previously described ^30.^

#### Image processing and imaging data analysis

All image processing was performed using Zeiss Zen 2.1 v11, Adobe Photoshop and Illustrator, NIH ImageJ, Bitplane Imaris 8.3, and custom Matlab scripts.

Computational tracing of nerve fibers to establish global innervation patterns was performed using a customized version of the open-source software neuTube ^66^ to perform initial batch tracing of image tiles. The resulting tracings were stitched, filtered, and visualized using custom Matlab scripts. Tracing of cholinergic neurons from intrinsic ganglion to the SA node was performed semi-automatically using neuTube.

Quantification of cell transduction in intrinsic cardiac ganglia (Figure 2c and Supplemental Figure 1e), stellate ganglia, and medulla sections (Supplemental Figure 1b) was performed by manual cell counting using NIH ImageJ Cell Counter.

For the stellate ganglion heat maps in Figure 5, 3D locations of CTB-labeled cell bodies in each stellate ganglion were marked using NIH ImageJ Cell Counter. Maximum intensity projection images were manually annotated with 2D polygonal regions of interest demarcating the extent of the stellate ganglion. Marked cell bodies outside the stellate ganglion were excluded from subsequent analysis. Landmark point locations were selected along the stellate ganglion boundary (e.g., ansa). One right and one left stellate ganglion image were selected to serve as reference shapes (white dashed outline in Figure 5d). To align the landmarks for each sample with the selected reference sample, a non-rigid warping was estimated by fitting a 2D thin plate spline warp ^67^ with a regularization parameter of 1e^-6^. The density of labeled cell bodies was estimated by binning the x,y-image coordinates of each marked cell into 50 × 50 µm bins and counting the number of cells falling in each bin in order to compute the number of cells per square µm in projection. These density maps from each individual sample were then warped using the thin plate spline warp estimated from the spatial landmark correspondences to align them all with the reference sample. Finally, the densities were averaged point-wise across samples. One could, in principle, perform warping and density estimation in 3D. However, we observed little variation in the densities labeled of cells along the optical z-axis, presumably due to the consistent orientation of samples on slides.

### Statistics

Data are presented as mean +/- s.e.m.. Sample sizes are indicated in figure legends or text. Comparisons for all pairs were conducted with a two-tailed Student’s t test. P < 0.05 was considered statistically significant.

## AUTHOR CONTRIBUTIONS

P.S.R., R.C.C., J.L.A., V.G., and K.S. designed the study. S.H. and P.S.R. performed heart clearing. A.G. and P.S.R. performed lightsheet imaging of cleared hearts. K.Y.C., B.E.D., R.C.C., and V.G. designed viral constructs. R.C.C. and K.Y.C. produced viruses. C.C.F. developed computational pipeline, performed fiber tracing, and analyzed stellate ganglia CTB data. P.S.R. and R.C.C. performed immunohistochemistry. P.S.R. performed optogenetic experiments. J.D.T. helped with optogenetic experiments. M.C.J. performed cardiac CTB injections. P.S.R., P.H., S.H., B.A.G-.W., and G.S. analyzed data. H.M. provided guidance on TH mouse line selection. P.S.R. prepared the figures. P.S.R. and K.S. wrote the manuscript. All coauthors contributed to the final version of the manuscript.

## ACKNOWLEDGEMENTS

We thank the entire Shivkumar and Gradinaru group for discussions. This work was supported by a NIH Stimulating Peripheral Activity to Relieve Conditions (SPARC) awards (OT2OD023848 to K.S. and V.G.; OT2OD23864 to H.M.). The K.S. laboratory is also supported by a NIH National Heart, Lung, and Blood Institute (NHLBI) grant (R01HL084261) and a NIH SPARC award (U01EB025138-01 to K.S. and J.L.A.). The V.G. laboratory is also supported by a NIH Director's New Innovator Award (DP2NS087949), a NIH Presidential Early Career Award for Scientists and Engineers, a NIH National Institute on Aging grant (R01AG047664), a NIH BRAIN Initiative award (U01NS090577), the Defense Advanced Research Projects Agency (DARPA) Biological Technologies Office, the National Science Foundation NeuroNex Technology Hub (1707316), the Curci Foundation, the Beckman Institute, and the Rosen Bioengineering Center at Caltech. V.G. is a Heritage Principal Investigator supported by the Heritage Medical Research Institute. P.S.R. was supported by a NIH NHLBI F31 Predoctoral Fellowship (F31HL127974). R.C.C. was supported by an American Heart Association Postdoctoral Fellowship (17POST33410404). P.S.R. is a part of the UCLA-Caltech Medical Scientist Training Program. We thank Drs. Marmar Vaseghi (UCLA), Olujimi A. Ajijola (UCLA), and Ching Zhu (UCLA) for their comments on the manuscript.

## Notes

The authors have declared that no conflict of interest exists.

